# Identification and characterization of interactions between Influenza A Virus NS1 protein and the human ubiquitin proteasome system

**DOI:** 10.1101/2023.10.05.561086

**Authors:** Laurie-Anne Lamotte, Samuel Kindylides, Chloé Gaupin, Caroline Demeret, Lionel Tafforeau

## Abstract

As a key player involved in various cellular pathways, including innate immune response activation, the human ubiquitin-proteasome system (UPS) is particularly targeted by viral proteins upon infection. Indeed, most viruses have evolved to counteract and hijack this system, as it is the case for the influenza A virus (IAV). The non-structural protein 1 (NS1) is described as the main IAV virulence factor, which is known to interact with several cellular proteins, including some UPS factors that are important for the viral escape of the immune cell response. In this study, we profiled the overall interplay between the NS1 proteins of multiple IAV strains and the human UPS. We identified 98 UPS factors directly interacting with NS1 of all or a subset of the studied strains, and we functionally studied 18 of them. We highlighted the involvement of these UPS factors in the IAV life cycle by performing viral titrations, minigenome replicon assays and an ISRE-luc (IFN pathway) assays. Furthermore, we analyzed the expression and subcellular localizations of FZR1, MKRN3, RC3H2 and SHKBP1 upon IAV infection. This interactomics approach allows for an improved understanding of the interplay between NS1 and UPS pathway in the context of an IAV-mediated inhibition of cellular anti-viral responses.

**Importance:** Influenza A viruses (IAV) are pathogens responsible for annual flu epidemics causing up to 650,000 deaths each year, resulting in a significant impact in public health and global economy. IAV are also responsible of occasionally pandemic outbreaks in human population, such as in 1918 that caused the death of 50-100 million people. Non-structural protein 1 (NS1) is the main IAV virulence factor; it acts by direct interactions with several cellular proteins, leading to the host shut-off and to the inhibition of the host cell innate immune response. Since the ubiquitin-proteasome system (UPS) plays a crucial role in the innate immune response activation, it is a designated target for NS1 upon infection. Our research thus focused on the identification of interactions between NS1 of 6 different IAV strains and the UPS, to better understand the interplay between this viral protein and the UPS upon viral infection.

## 1. Introduction

Ubiquitination is a post-translational modification that regulates the fate of target proteins by modifying their stability, localization, and activity (1, 2). Conjugation of an ubiquitin molecule (Ub) on a specific substrate requires a sequential three-step enzymatic cascade involving E1, E2 and E3 enzymes. Ub is first bound to an E1 ubiquitin-activating protein, in an ATP-dependent way. It is then transferred to an E2 ubiquitin-conjugating enzyme. Finally, an E3 ubiquitin-ligase binds ubiquitin to a lysine (K) on the protein substrate (3–6). Substrates can then either be mono-ubiquitinated or poly-ubiquitinated, through the formation of ubiquitin chains via internal lysine residues within ubiquitin itself. The best described poly-ubiquitin chains are K48-linked chains, which are mostly associated with protein degradation by the 26S proteasome, and K63-linked chains, which are involved in trafficking, endocytosis, and signal transduction (7–9). Ubiquitinated substrates can subsequently be recognized by ubiquitin-binding domain-containing proteins, such as de-ubiquitinating enzymes (DUBs). DUBs are able to hydrolyze isopeptide bonds formed between ubiquitin moieties, or between ubiquitin and the protein substrate (10, 11). The whole ubiquitination system, including E1, E2, E3 and DUB enzymes, as well as the 26S proteasome, is commonly referred to as the ubiquitin-proteasome system (UPS). Depending on the type of ubiquitination, the substrate protein will undergo different fates. Because of its crucial role in innate immune response activation, notably through the activity of TRIM E3 ligases, the UPS is a designated target for viral proteins upon infection (12, 13). As an example, TRIM25, whose activity consists in promoting RIG-I sensor activation and subsequent innate immune response signaling, was described to be particularly inhibited by the influenza A virus (IAV) non-structural protein 1 (NS1) (14, 15).

Influenza A virus is a member of the *Orthomyxoviridae* family, and its genome consists of eight single-stranded negative sense RNA segments that encode different proteins. The hemagglutinin (HA) and neuraminidase (NA) glycoproteins are found at the surface of the viral particle, together with the M2 ion channel protein. The viral envelope itself surrounds a matrix made of M1 protein oligomers, that encompasses the segmented genome. This genome consists of RNA segments encapsidated by many nucleoprotein monomers (NP) and associated with an RNA-dependent RNA polymerase (PB1, PB2 and PA subunits). The genome also encodes, among other proteins, a nuclear export protein (NEP, also named NS2) and the forementioned NS1 protein (16). NS1 is described as the main IAV virulence factor, acting by direct interactions with cellular proteins or through the regulation of host genes expression, a process named host shut-off (17–25). In addition to TRIM25, some other perturbations of UPS factors by NS1 were described, such as the regulation of the MDM2 E3 ligase, OTUB1 DUB, and A20 E3 ligase/DUB (26–30).

More generally, the interplay of viral proteins with the UPS may impact on viral evolution or pathogenic power, as it has been shown previously for the human papillomavirus proteins (31, 32), as well as for the IAV PB2 protein (33). In this study, we sought to delineate the interplay between the UPS and the NS1 protein of IAV strains of various virulence. We thus screened an human UPS-dedicated library encoding for about half of the whole human UPS for interaction with NS1 proteins of six viral strains, using the *Gaussia princeps* protein complementation assay (GPCA). From these screenings, we then selected some interacting UPS factors in order to study their functional implication in IAV life cycle. Upon performing virus titration assays in siRNA-depleted cells, we showed that several UPS factors are involved in proper viral replication. Furthermore, we observed that several UPS factors are necessary to activate the interferon pathway in infected cells. Finally, we analyzed the expression and subcellular localization of a subset of 4 UPS proteins upon IAV infection, namely FZR1, MKRN3, RC3H2 and SHKBP1.

## 2. Results

### 2.1. Identification of UPS factors interacting with IAV NS1 protein

In order to perform a comparative interactomics analysis between the UPS and IAV NS1 proteins, we conducted a GPCA screening using the whole UPS library, containing 737 cDNAs encoding for 585 different UPS factors, and NS1 from six different IAV strains. Four human influenza strains were selected: H1N1_WSN_, H1N1_1918_, H1N1_pdm09_, and H3N2, and one avian zoonotic H7N9 strain, that had (H7N9, isolated from an infected individual) or had not (avian H7N9, isolated from an infected poultry) passed the human species barrier.

The NS1-UPS screening was conducted by co-expressing in H2K293T each of the NS1 protein fused with the C-ter hemi-Gaussia luciferase and each UPS expressed in fusion with the N-ter hemi-Gaussia luciferase. In such matrix-based screening, the 6 NS1 proteins were assessed for interactions with the same set of UPS factors in the same experiment, in order to ensure strict comparative results.

For each NS1 screen, a positive threshold, based on the distribution of the luminescence values (RLU), was calculated as the third quartile + 1.5 times the interquartile range (Q3 + 1.5 IQR). UPS factors-NS1 pairs generating luminescence values above this threshold were considered as potential positive direct interactions. Of the 737 UPS factors tested, 132 were found to interact with the NS1 protein from either one or more strains (Figure S1, Table S1). To note, the mean of luminescence values varies for each NS1, with higher values for H1N1_pdm09_ (mean RLU= 3,084,216), and lower values for H3N2 (mean RLU= 100,154). This was correlated with the expression level of each NS1 protein, observed through western blot (Figure S2). The UPS interactors list was then cleared from redundant ORFs or known sticky factors (such as NEURL, UBA, UBB and USP35) to finally obtain 112 potential interactors.

These 112 UPS factors were then retested against the 6 NS1 proteins through a second GPCA screening, by applying the normalized luminescence ratio method (NLR) (37). In this secondary screen, the NS1 proteins were also tested against a random reference set (RRS) of 20 proteins *a priori* not interacting with NS1, used to set the threshold of positive interactions (see methods). Such NLR-based protein-protein interaction (PPI) profiling takes into account the interaction background of each partner and has already been shown to accurately discriminate interacting partners from false positives (33, 37). Of the 112 retested UPS factors, 98 were validated as interacting partners of at least one NS1 protein (Figure 1A, Table S2). Interestingly, among these positive candidates, 22 are common with the 6 NS1 tested, and 17 interactions are specific to NS1 from one strain (Figure 1B).

**Figure 1:**
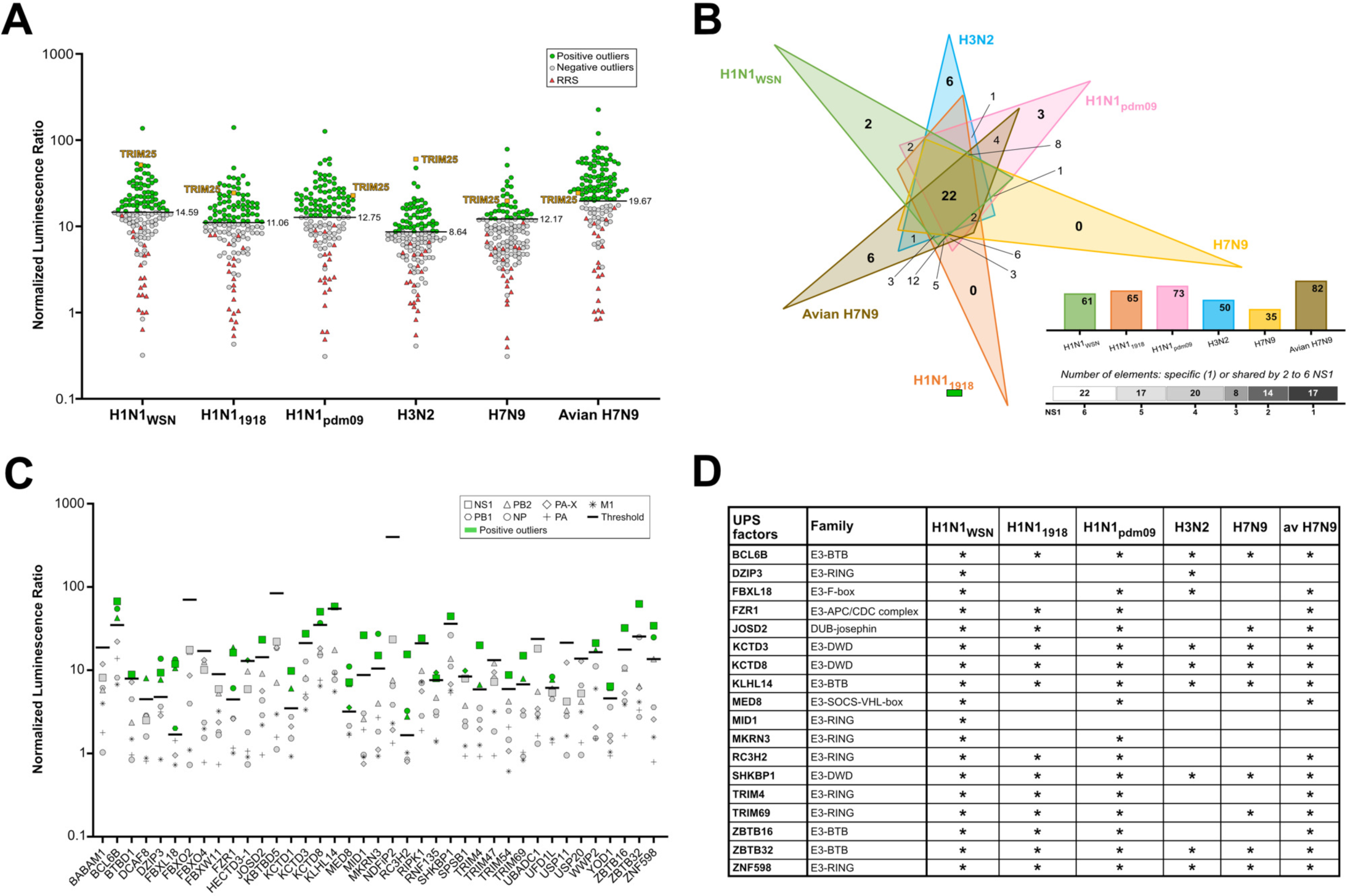
IAV NS1 interacts with the human UPS. **A.** Results of the second GPCA screening performed between 112 UPS factors identified in the first GPCA screening, and NS1 from H1N1_WSN_, H1N1_1918_, H1N1_pdm09_, H3N2, H7N9 and avian H7N9 strains. Normalized luminescence ratio (NLR) was calculated for each protein pair. The positive threshold (black bold lines) was calculated for each NS1 protein from NLR obtained from random reference set (RRS). NLR of RRS are represented as red triangles, NLR of negative putative interactors as gray circles and NLR of positive interactors as green circles. As an example of positive interactors, TRIM25 is depicted as yellow squares. **B.** Venn diagram and tables indicating the number of UPS factors interacting with each NS1 protein and numbers of specific or shared interactions. **C.** Results of the third GPCA screening conducted between 39 positive UPS factors and H1N1_WSN_ viral proteins. NLR, and positive threshold (black bold lines) were calculated for each UPS factor. Values for viral proteins are represented in gray (different symbol for each viral protein) and positive viral proteins outliers are shown in green. **D.** Table summarizing the selected 18 UPS factors and their respective UPS families. Their interactions with NS1 from the 6 different IAV strains are represented by a star. UPS factors families are based on iUUCD 2.0 classification (63).

A literature-based curation was carried out to keep only the positive factors with a reported function in protein ubiquitination, since the UPS library was gathered based on the presence of peptides motif characteristics of different categories of UPS factors (32), to a final selection of 39 recognized interactors. These UPS factors were screened against NS1 and other H1N1_WSN_ IAV proteins, as well as against 20 RRS. A threshold was calculated for each Glc1-UPS, based on NLR from RRS tested in GPCA. We notably observed that two RRS proteins were particularly sticky (RRS8 GSTT1 and RRS18 TRPT), depicted by their high NLR values, resulting in particularly high, potentially over estimated, threshold for positive viral proteins/UPS PPIs (Table S3). Conversely, FBXO2, KBTBD5 and NDFIP2 UPS factors strongly bind to some factors of the RRS, which leads to their respective high threshold values and may have blurred their interaction with NS1 (Figure 1C, Table S3). In such settings, 25 UPS factors were validated as NS1 interactors, which represent top high-confidence NS1 interactors (Figure 1C, Table S3). We noticed that, among the UPS interacting with NS1, several are also interacting with other IAV proteins (mostly PB2 and NP) (Figure 1C, Table S3).

### 2.2. Functional impact of UPS factors on IAV replication

To determine the biological relevance of identified UPS factors and NS1 protein interplays, we examined the implication of NS1 interactors in IAV replication cycle, of which 18 were selected for further functional characterization (Figure 1D, Table S3). The production of infectious viral particles upon siRNA-mediated depletion of eah NS1 interactor was assessed by the plaque assay method. Since the depletion of the UPS factors ATXN3 (Josephin DUB), RNF40 and TRAF2 (RING E3 ligases) was previously shown to significantly impair H1N1_WSN_ virus replication, these 3 UPS factors were used as controls (33). A549 cells, transfected with a non-targeting siRNA (SCR) or with siRNAs targeting the individual UPS factors or the three controls (2 siRNAs/factor), were infected for 24 hours with H1N1_WSN_ virus at a MOI of 10^−4^. Knock-down efficiency and cell viability (Figure S3) were determined for each siRNA. The effect of UPS factor depletions was calculated by comparing viral titer obtained in siRNA conditions with that of the SCR control.

Of the 18 UPS factors, 12 were shown to have an impact on viral replication. Both siRNAs used for FZR1, RC3H2 and ZNF598 depletion significantly impaired IAV replication, with 0.18-0.28-fold, 0.33-0.43-fold and 0.44-0.74-fold decreases, respectively, suggesting that these proteins may act as pro-viral proteins (Figure 2, Table S6). Several other UPS factors have significant reducing effects on viral production with only one siRNA. On the contrary, some UPS factors knock-downs were shown to significantly promote viral replication, such as DZIP3, KCTD3 and ZBTB32 (p value <0.05).

**Figure 2:**
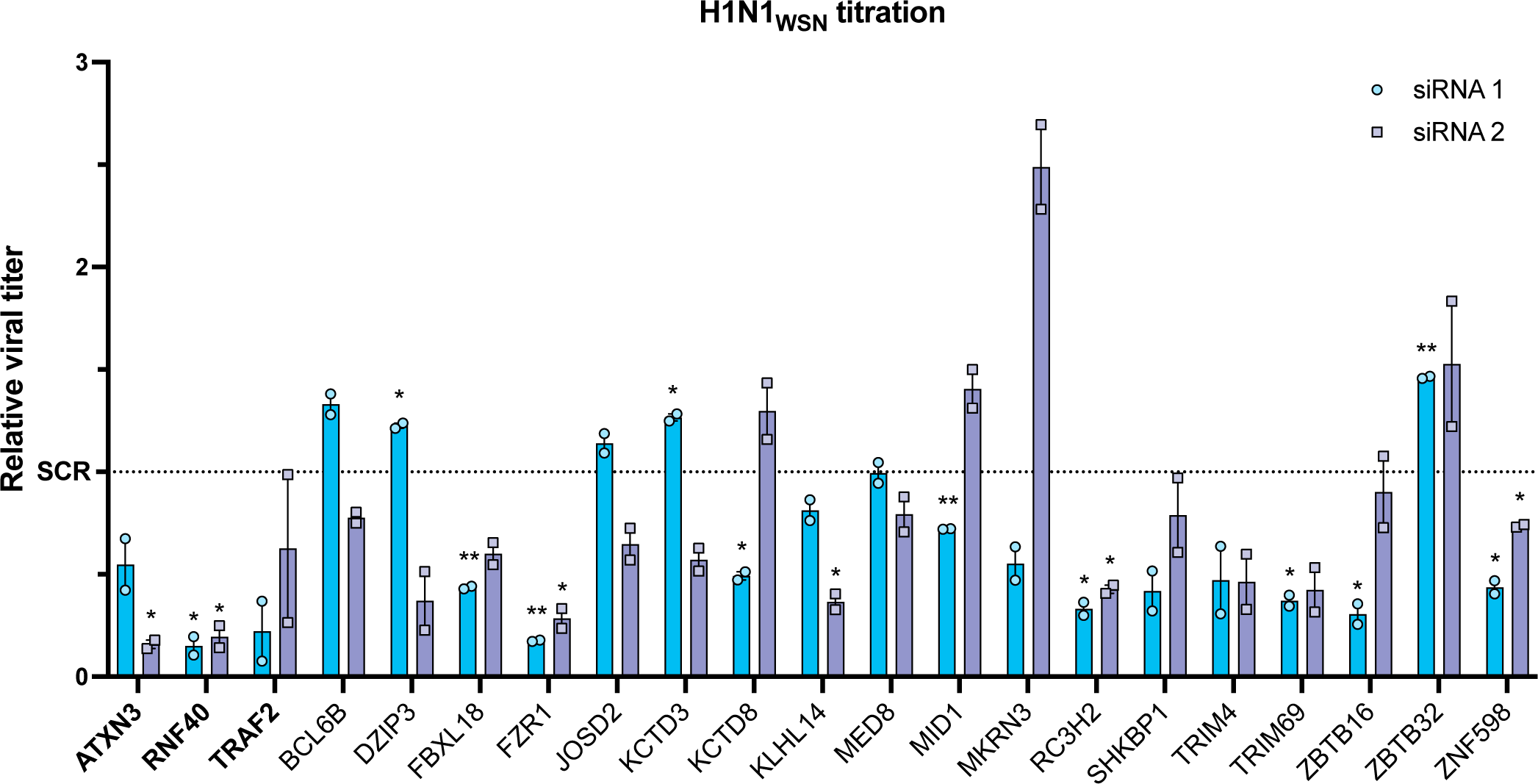
Effects of UPS factors depletion on H1N1_WSN_ virus replication. A549 cells were transfected with UPS factor targeting-siRNAs (2 siRNAs/factor) for 48h and were infected for 24 hours with H1N1_WSN_ virus at a MOI of 10^−4^. Results of the plaque assay are represented as relative viral titer, obtained by comparing viral titers obtained in siRNA transfected cells to SCR transfected control cells. Data represent the mean ± SD of two independent experiments performed in triplicates, and p values were calculated with unpaired t-tests, *<0.05, **<0.01. ATXN3, RNF40 and TRFA2, in bold, are literature controls. SCR: non-targeting siRNA.

To determine the effect of the UPS factors during the course of a single viral replication cycle, a titration assay based on a Nano-luciferase reporter virus was performed. A549 cells were transfected with siRNAs and then infected with a H1N1_WSN_ virus carrying the Nano-luciferase ORF in its PB2 segment (36). In this assay, luminescence values are directly correlated to viral titers, as it was demonstrated through a standardization experiment (Figure S4). Viral particles were collected during a virus replication cycle (after 3 hours, 6 hours, 9 hours, and 12 hours of infection), then used to infect for 6h non-treated A549 cells, and luminescence values obtained in each condition were compared to SCR-transfected cells. Among the 18 UPS factors, the depletion of 7 of them showed no impact on viral replication, independent of the time point observed (Figure 3). For the other 11 factors, their depletion modified viral replication compared to the control, mostly at 3 hours and 12 hours post-infection (hpi). Strikingly, UPS factors depletion positively impacted IAV replication at the beginning of the viral cycle, which is notably the case for JOSD2, MED8, MKRN3 and RC3H2. At late stages of the viral replication cycle (9 - 12 hrs), both positive and negative effects were observed . The most significant impact of UPS factors depletion appeared for ZNF598 and FZR1, for which the 2 siRNAs decreased viral replication at 12hr, albeit to different extent (Figure 3). We noticed an effect of ZBTB16 siRNA 1 which should be due to its toxicity but not siRNA depletion (Figure S3B).

**Figure 3:**
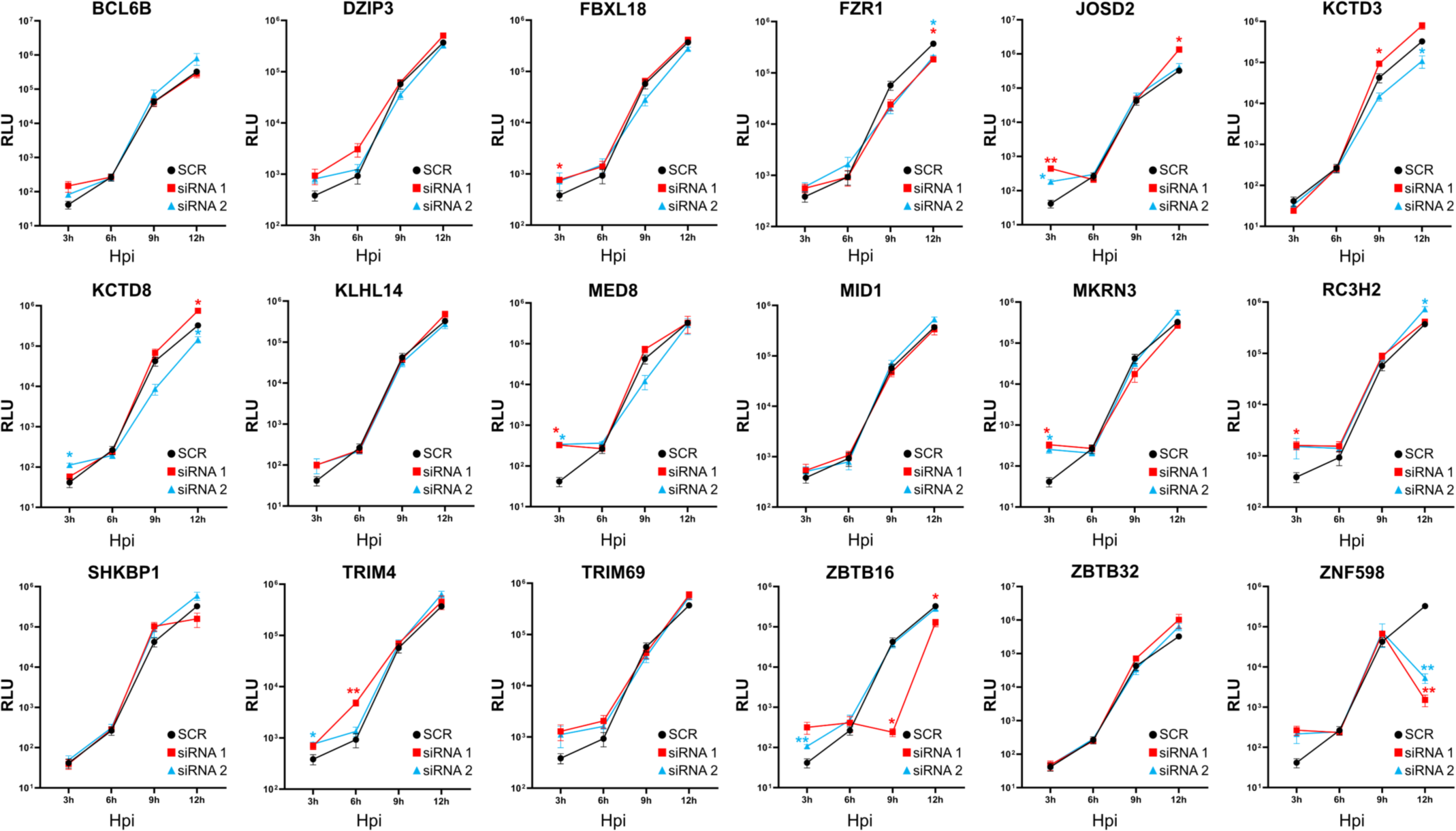
Effects of UPS factors depletion on IAV replication during the course of infection. A549 cells were transfected with siRNAs (2 siRNAs per UPS factor) for 48h and were then infected with H1N1_WSN_ PB2-Nanoluc reporter virus at a MOI of 10^−4^ for 3 hours, 6 hours, 9 hours, and 12 hours. Results are expressed as relative luminescence units (RLU). For each UPS factor, SCR luminescence values are shown in black, siRNA 1 values in red and siRNA 2 values in blue. Data represent the mean ± SD of four independent experiments, performed in triplicates, and p values were calculated by two-way ANOVA followed by Dunnett’s multiple comparisons test, *<0.05, **<0.01. Hpi: hours post-infection. SCR: non-targeting siRNA.

### 2.3. Functional impact of UPS factors on viral polymerase activity

To investigate whether these 18 UPS factors may alter IAV polymerase activity, we performed a minigenome replicon assay. Almost all knock-down conditions led to statistically significant modifications in viral polymerase activity, except for ZNF598, even though these modifications were mostly minor compared to the control SCR (Figure 4). However, several UPS factor depletions stood out from the rest. The depletion of KCTD8, MKRN3, TRIM69 and ZBTB16 led to a higher polymerase activity, ranging from 1.2 to 2.57-fold increase, but with only one of the two siRNAs employed. On the contrary, BCL6B, DZIP3, JOSD2, MID1, RC3H2, TRIM4 and ZBTB32 depletion with each siRNA reduces the polymerase activity, ranging up to 3,8-fold decrease for TRIM4 siRNA 1 (Figure 4).

**Figure 4:**
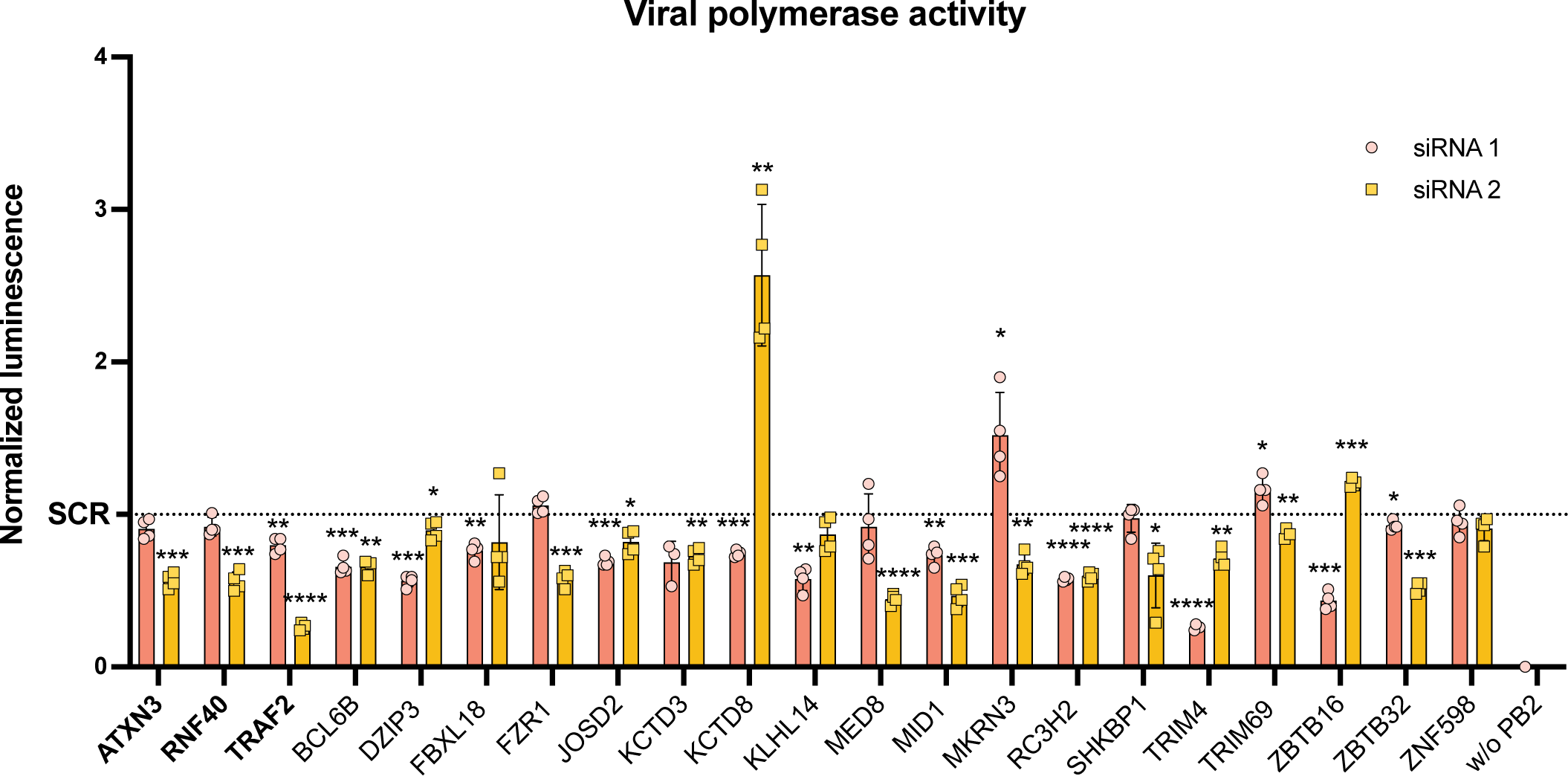
Effects of UPS factors depletion on viral polymerase activity. Each UPS factor was first depleted by the transfection of 2 siRNAs in HEK-293T cells. Cells were then transfected with a set of plasmids coding for viral proteins (PB1, PB2, PA, NP, and NS), with a plasmid coding for the firefly luciferase, as a reporter, and with a plasmid coding for the *Renilla* luciferase, as a transfection efficiency control. Luminescence values obtained with firefly luciferase were normalized to values obtained with *Renilla* luciferase, and values obtained for siRNA conditions were then normalized to values obtained with SCR. Cells transfected with the complete set of plasmids, but lacking the PB2 plasmid, served as control for plasmid set efficiency (w/o PB2). Data represent the mean ± SD of four independent experiments, and p values were calculated with unpaired t-tests, *<0.05, **<0.01, ***<0.001, ****<0.0001. ATXN3, RNF40 and TRFA2, in bold, are literature controls. SCR: non-targeting siRNA.

### 2.4. Functional impact of UPS factors on IFN pathway

As IAV infection is known to subvert host innate immune response, notably through perturbations caused by NS1 protein, we aimed to determine if the 18 UPS factors may be involved in IFN response. We previously generated an A549 cell line which expresses the firefly luciferase upon IFN stimulation, by the presence of ISRE upstream of a minimal promoter. We demonstrated that this A549-RING1 cell line properly responds to IAV infection (34). We therefore treated these A549-RING1 cells with siRNAs targeting the 18 UPS factors, to determine the individual impact of each UPS factor on the IFN pathway (Figure 5A). Of the 18 UPS factor depletions, 8 induced a higher luminescence signal compared to control cells (DZIP3, FBXL18, FZR1, KLHL14, MKRN3, RC3H2, TRIM4 and TRIM69) and 7 decreased this signal (BCL6B, JOSD2, KCTD8, MED8, SHKBP1, ZBTB16 and ZNF598).

**Figure 5:**
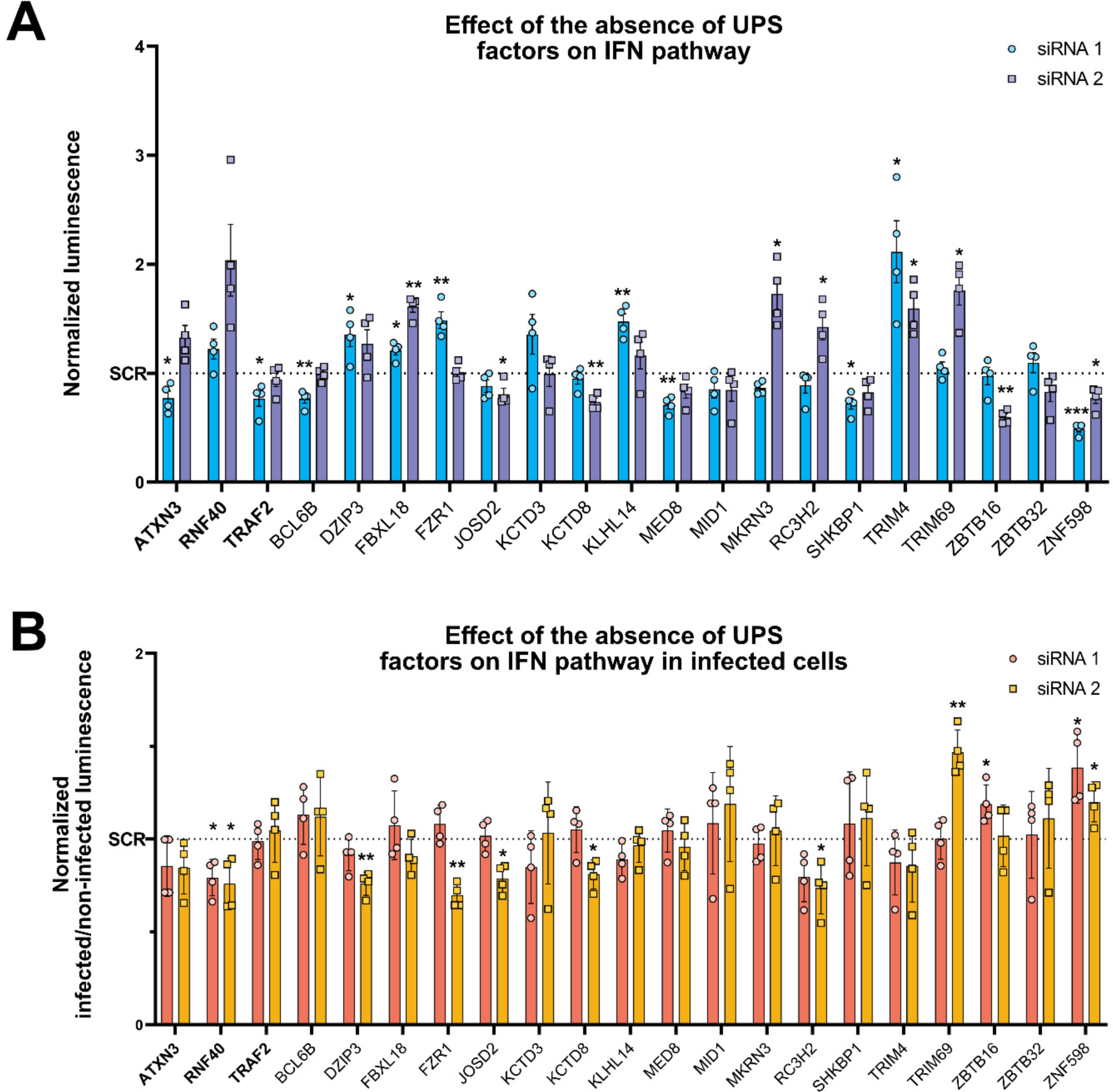
Effects of UPS factors depletion on IFN pathway in non-infected and infected cells. A549-RING1 cells, which express firefly luciferase upon ISRE minimal promoter, were transfected with siRNA targeting the 18 UPS factors (2 siRNAs per factor). Cells were then either left non-infected (A) or were infected with H1N1_WSN_ virus at a MOI of 10^−4^ (B) for 48 hours. A. Luminescence values obtained in non-infected cells were normalized to luminescence values obtained in SCR control, for each condition. B. Luminescence values obtained in infected cells were normalized to luminescence values obtained in SCR control, and then normalized to values obtained from non-infected cells as given in (A). Data represent the mean ± SD of four independent experiments, and p values were calculated with unpaired t-tests, *<0.05, **<0.01, ***<0.001. ATXN3, RNF40 and TRFA2, in bold, are literature controls. SCR: non-targeting siRNA.

Next, we infected these UPS factor-depleted A549-RING1 cells with H1N1_WSN_ for 48 hours at a MOI of 10^−4^ (Figure 5B). We observed that the depletion of 8 UPS factors (DZIP3, FZR1, JOSD2, KCTD8, RC3H2, TRIM69, ZBTB16 and ZNF598) induced significant changes in IFN response upon infection as compared to non-infected cell. Notably, the depletion of RC3H2 decreased luciferase expression under IAV infection, and thus IFN production, suggesting that this UPS factor may be involved in host anti-viral response. Conversely, ZNF598 knock-down induced a higher luciferase expression as compared to non-infected cells, with 1.38 and 1.2-fold changes (respectively for siRNA 1 and 2). Another important luciferase expression modification came from TRIM69-depleted cells (siRNA 2) upon IAV infection, which increased luminescence values to 1.47-fold.

### 2.5. Expression and subcellular localization of selected UPS factors upon IAV infection

The IAV NS1 protein is known to shuttle between the nucleus and the cytoplasm of infected cells, through its nuclear export signal and its two nuclear localization signals, of which one may also act as nucleolar localization signal (42, 43). We therefore wondered if NS1 may change UPS factor subcellular localization during the course of infection to modify, for instance, their activity or stability. To answer this question, we selected four UPS factors of interest, based on forementioned results: (i) FZR1, also named Cdh1, whose depletion decreased viral replication, while its transcription was increased upon infection (Figure S5); (ii) MKRN3, whose knock-down impaired IAV replication at 3 hpi and modified viral polymerase activity; (iii) RC3H2, whose depletion significantly decreased polymerase activity and IFN pathway response upon infection; (iv) and SHKBP1, one of the 22 common NS1 interactors and which only interacted with NS1 among the multiple IAV proteins tested. A549 cells were infected with H1N1_WSN_ and the subcellular localization of these UPS factors, as well as NS1, was monitored during the virus infection cycle (0h, 3h, 6h, 9h and 12hpi). As expected (44), NS1 localization changes during the viral life cycle, beginning with a cytoplasmic localization (3-6 hpi), followed by a major relocalization in the nucleus and nucleolus (9 hpi). At 12 hpi, undoubtedly for a multi-cycle infection too, NS1 is highly expressed in the cytoplasm, in the nucleus and in the nucleolus at the same time (Figure 6 A-D). Of the 4 UPS factors, only RC3H2 did not change localization along infection, even though RC3H2 and NS1 share the same cytoplasmic localization at 6 and 9 hpi (Figure 6D). Before IAV infection, FZR1 was clearly expressed in the cytoplasm as elongated structures, and then began to localize in cytoplasmic foci in infected cells (3 hpi). These foci accumulated at 6 hpi and at 9 hpi and were interestingly devoid of NS1 (white arrows at 6 and 9 hpi). During the later stages of infection, these foci localizations decreased, and FZR1 mostly regained its elongated aspect (Figure 6A). In mock-infected cells, MKRN3 was expressed in the cytoplasm, as well as in the nucleus, with a clear nucleolus exclusion. During the early stages of IAV infection (3 hpi), MKRN3 was excluded from the nucleus and co-localized with NS1 in the cytoplasm. At 6 hours post-infection, MKRN3 took the shape of little dots notably organized around the nucleus. Following NS1 entry in the nucleus and nucleolus at 9 hpi, MKRN3 accumulated in the nucleus and was less localized in the peri-nuclear regions rich in NS1 (white arrows at 9 hpi). This exclusive localization was still observable at 12 hpi, while MKRN3 recovered its original localization (Figure 6B). Concerning SHKBP1, originally localized at cell junctions and in the nucleus (white arrows at 0 hpi), it began to localize in the cytoplasm at 3 hpi. At 6 hours post-infection, SHKBP1 traded its cytoplasmic and cell junction localizations for a high nuclear accumulation, only visible in cells in which NS1 is nucleolar (white arrows at 6 hpi). This change in localization was reversed at 9 hpi, when SHKBP1 regained cell junctions (white arrows at 9 and 12 hpi), while NS1 was still expressed in nucleoli (Figure 6C).

**Figure 6:**
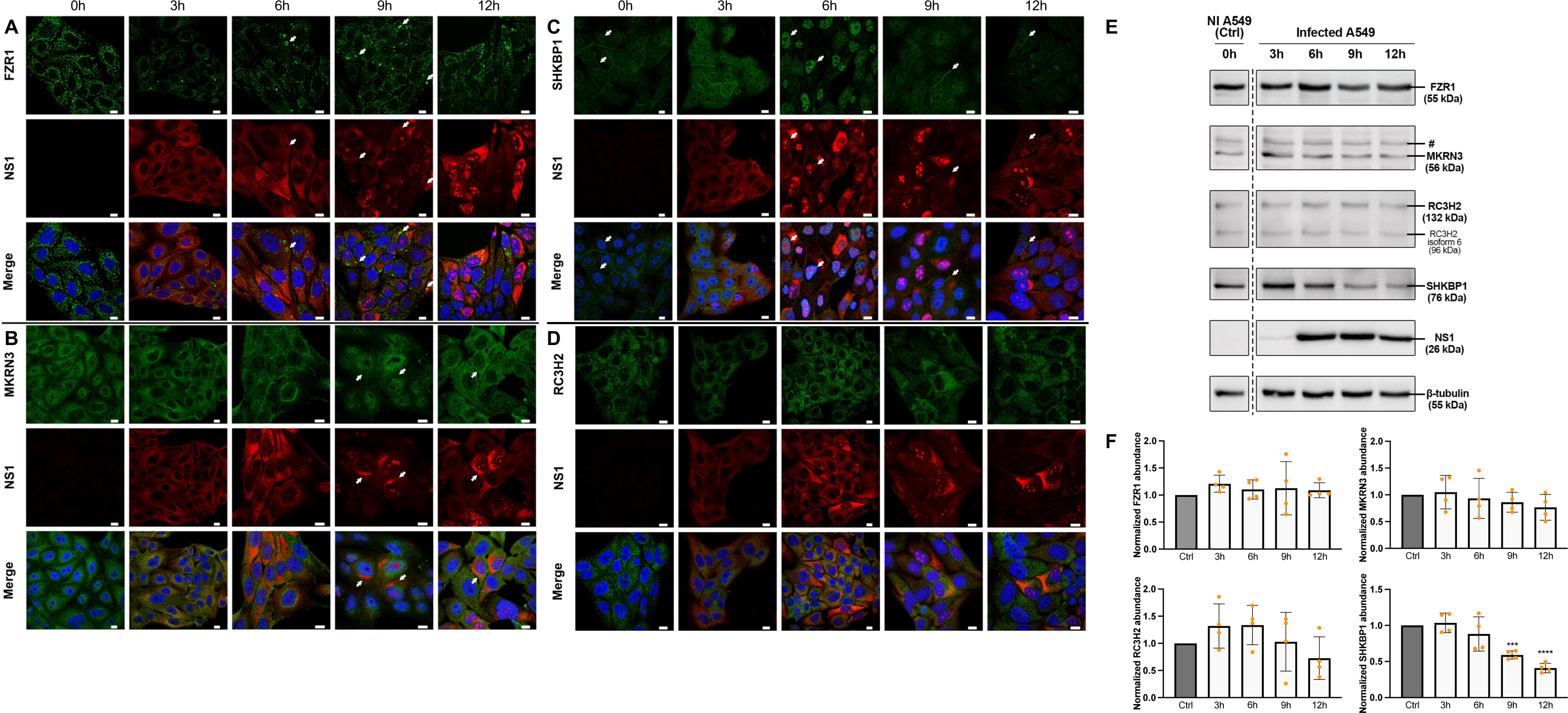
FZR1, MKRN3, SHKBP1 and RC3H2 localization and protein level during the course of IAV infection. A549 cells were infected with H1N1_WSN_ virus at a MOI of 3. **A-D.** Cells were fixed at different time points of infection (0, 3, 6, 9 and 12 hours) and subjected to immunofluorescence with antibodies targeting endogenous FZR1 (**A**), MKRN3 (**B**), SHKBP1 (**C**), RC3H2 (D) and NS1. Most interesting localizations are highlighted with white arrows. Green: UPS factors. Red: NS1 protein. Blue: DAPI staining of nuclei. Scale bars: 10 µm. E-F. Proteins were extracted at different time points of infection (0, 3, 6, 9 and 12 hours) and subjected to western-blot. (**E**) representative western-blots of the selected UPS factors. # represents a non-relevant cross-reaction band revealed with MKRN3 antibody. (F) Densitometric analyses of protein expression were performed with ImageJ and normalized to ctrl (non-infected cells). Data represent the mean ± SD of four independent experiments, and p values were calculated with unpaired t-tests, ***<0.001, ****<0.0001.

The protein abundance of the four UPS factors was also assessed along the virus infection cycle (0h, 3h, 6h, 9h and 12hpi), as well as NS1, as a positive control of infection. IAV infection did not affect FZR1 expression, that remained stable throughout the infection. On the contrary, we observed a decrease of MKRN3, RC3H2 and SHKBP1 abundance during the late stages of infection, with a significative reduction of SHKBP1 at 9 and 12 hpi (Figure 6E-F).

## 3. Discussion

Given the cellular implications of the ubiquitin-proteasome system (UPS), notably in innate immune response regulation, it is not surprising that most viruses evolved to subvert or exploit the UPS pathway (45). One of the best described interactions found between the UPS and a viral protein is the TRIM25 E3 ligase and the influenza A virus (IAV) NS1 protein interplay (15). As NS1 is the main IAV virulence factor, it may undoubtedly interact with other UPS factors (30). In this study, we therefore first identified the interplay between NS1 and the human UPS. Next, we characterized the functional role of these interactions in the IAV life cycle.

Through two successive *Gaussia princeps* protein complementation assay (GPCA) screenings, we showed that the NS1 protein interacts with 98 UPS factors out of the 737 listed in the GPCA library (Figures 1A and S1). Interestingly, 22 interactions are common between the 6 NS1 proteins tested, all coming from different IAV strains, suggesting an evolutionary conserved involvement of these UPS factors in IAV infection (Figure 1B). Other interactions appear to be more strain-specific. This may be explained by the various degree of virulence of the employed strains, i.e., laboratory-adapted H1N1_WSN_ strain, slightly virulent seasonal circulating IAV strains (H1N1_pdm09_ and H3N2), highly virulent H1N1_1918_ and H7N9 strains, as well as avian H7N9 strain.

Following a literature-based review, 39 UPS factors were selected and assessed for interaction with the other H1N1_WSN_ proteins (except the surface glycoproteins HA and NA and the ion channel protein M2). 25 of them were recovered as NS1 interactors, but also with other IAV proteins, mainly PB2 and NP (Figure 1B, Table S3). The interaction redundancy between several viral proteins and those cellular factors may underline a combined regulation of the UPS to orchestrate the viral infection cycle, such as in host shut-off, cap-snatching or innate immunity evasion, that are not exclusively relying on NS1. Finally, 18 UPS factors, found to be mostly E3 ligases, were selected for further characterization (Figure 1D). This stringent selection allowed us to only study the functional impact of UPS factors which displayed a strong NS1 interaction (correlated to GPCA signals), even though it did not invalidate other interactions found through the second GPCA screening.

We first investigated the role of these 18 UPS factors in H1N1_WSN_ viral replication by performing a plaque titration assay (39). We showed that the depletion of FZR1, RC3H2 and ZNF598 impaired IAV replication (Figure 2). We then monitored the effect of UPS factor depletions on virus production throughout a single replication cycle. Some effects of UPS factor knock-down at late stages of infection were found to be correlated with results obtained by plaque titration, performed 24 hours after infection. Indeed, FZR1 and ZNF598 depletion significantly impaired viral particles production at 12 hpi (Figure 3).

Minigenome replicon assay results showed that, apart from ZNF598, all tested UPS factors may have a slight impact on IAV polymerase activity, especially BCL6B, KCTD8, MID1, MKRN3, RC3H2, TRIM4, TRIM69 and ZBTB16 (Figure 4). However, their involvement in polymerase activity may not be linked to their interaction with NS1, as most of them were found to also interact with PB2 and/or NP proteins. Indeed, of all UPS factors tested, only JOSD2, KCTD3, MID1, SHKBP1, ZBTB16 and ZBTB32 were specific NS1 interactors, whose implication in viral polymerase activity may be related to NS1 (Figure 1C, Table S3). Moreover, it is important to note that IAV polymerase activity gathers both transcription and replication processes, which are not dissociated in minigenome replicon assay. In this study, we can therefore not assess if the slight effects of the UPS factors on viral polymerase activity are linked to either one or both phenomena. Nevertheless, effects of these UPS factors on viral polymerase activity are not reflected in viral replication, therefore suggesting another implication in viral life cycle that remains to be elucidated.

We generated an A549 ISRE-Luciferase cell line (34) and studied the implication of the 18 selected UPS factors on IFN pathway, in non-infected and in infected cells (Figure 5). Among all the results obtained in non-infected cells, some can be related to previous studies. The absence of BCL6B induces a diminution of luciferase expression, indicating an impairment of the IFN pathway. Indeed, BCL6B, also named ZBTB28, was recently shown to activate IFNARs, the type I IFN membrane receptors, thus enhancing the IFN transduction pathway (46). Such results are also observed upon ZBTB16 depletion, whose siRNA 2 induces a 0.6-fold diminution of luciferase expression. Upon IFN stimulation, ZBTB16 associates with PML and the histone deacetylase 1 to induce a subset of specific ISGs (47). As confirmed in our results, ZBTB16 is therefore directly involved in IFN response.

Of the 8 UPS factors whose depletion induces change in IFN response upon infection, some are particularly interesting. The second RC3H2 siRNA leads to a 26% decrease in innate immune response as compared to non-infected cells. RC3H2 was shown to promote ubiquitination and the subsequent degradation of apoptosis signal-regulating kinase 1 (ASK1), which promotes reactive oxygen species (ROS)-induced cell death (48). Among various biological functions, ROS are notably shown to promote the production of pro-inflammatory cytokines through the activation of NLRP3 inflammasome, a cellular complex itself activated by RIG-I (49, 50). This may in part explain the involvement of RC3H2 in IFN response upon infection that we depicted here. Concerning ZNF598, its knock-down promoted an increase of 1.2 to 1.38-fold in IFN response in the presence of the virus. Recent studies showed that ZNF598 is involved in innate immune responses during viral infection, through two modes of action. First, ZNF598 is involved in Ribosome-associated Quality control (RQC), which leads to ribosomal subunits dissociation and peptide degradation when ribosomes are collided (51). Through RQC, ZNF598 has recently been shown to negatively regulate ISG expression (52). Second, ZNF598 directly delivers a ubiquitin-like molecule, FAT10, to RIG-I, therefore impairing the IFN signaling cascade activation (53). In the absence of ZNF598 in IAV-infected cells, RIG-I should thus be more active and IFN response appears logically higher than in control cells. Consistent with this observation, innate immune response activation in ZNF598-depleted cells probably leads to the impaired IAV replication observed in the viral titration assays. ZNF598 transcription was also found to be highly increased upon IAV infection (Figure S5). NS1 may therefore interact with ZNF598, a designated pro-viral E3 ligase, in order to promote RIG-I inhibition and innate immunity evasion.

We also showed that protein expression and subcellular localization of selected UPS factors changes during the course of IAV infection (Figure 6). For example, SHKBP1, which is normally localized in the plasma membrane, where it is involved in the retention of epidermal growth factor receptor (EGFR) (54), undergoes a high nuclear accumulation at 6 hpi. By its presence in the nucleus, SHKBP1 is thus unable to prevent EGFR degradation, leading to the activation of the EGFR signaling pathway, which was found to promote IAV uptake (55). This pathway was also shown to promote G1/S transition (56). At an advanced infection stage, this SHKBP1 nuclear re-localization, which coincides with NS1 nucleolar localization, may be developed by the virus to limit the entry of new virions in host cells and to regulate the cell cycle. Moreover, we observed a significative decrease of SHKBP1 in infected cells. Intriguingly, FZR1 should normally be mainly expressed in the nucleus, as a component of the APC/C complex. However, we showed here that this protein is predominantly localized in the cytoplasm of A549 cells. This localization may be explained by the existence of two FZR1 isoforms (Cdh1α and Cdh1β), of which Cdh1β is found to be cytoplasmic, and notably expressed in A549 cells (57). Another explanation of this re-localization may be that FZR1 is phosphorylated in its nuclear localization signal region, thus regulating FZR1 activity through its cytoplasmic re-localization, resulting in G1/S transition (58). To note, the hijacking of cell cycle regulation machinery was already shown to be beneficial for IAV. Indeed, an arrest in G0/G1 phase was observed upon IAV infection and may be in part due to NS1 (59). Jiang *et al*. showed that NS1 inhibits the expression of both RhoA GTPase and cyclin D1, two crucial proteins for G1/S transition (60). Interestingly, FZR1 is also known to mediate the proteasomal degradation of RacGAP1, an important regulator of RhoA, while DZIP3 was found to stabilize cyclin D1 through mRNA interaction and K63-linked poly-ubiquitination (61, 62). The steady-state expression of FZR1 protein may thus be selected by IAV to regulate the cell cycle and to promote its replication. This hypothesis is also backed by the impairment of the production of viral particles observed in FZR1-depleted cells and the increase in FZR1 transcription upon infection (Figure S5). However, the localization of FZR1 in cytoplasmic foci, while its activity in G0/G1 phase maintenance seems crucial for IAV, and the absence of a co-localization signal with NS1, despite their interaction, remains to be elucidated.

In conclusion, we were able to identify UPS factors interacting with the NS1 protein and to perform a functional study of 18 selected factors. As these interactions are either conserved through IAV evolution or more strain-specific, the study of NS1 and UPS factors interplays taking into account strain virulence may be an interesting future approach. Among these NS1 interactors, some of them showed promising implications, notably in the IFN pathway, as well as some surprising re-localizations in infected cells. Nevertheless, more investigations are needed to decipher the role of these UPS factors in IAV life cycle.

## 4. Materials and Methods

### 4.1. Cells and viruses

Human embryonic kidney HEK-293T cells (ATCC #CRL-11268), human lung carcinoma cell line A549 (ATCC #CCL-185), stable cell line A549-RING1 (34), and MDCK-SIAT (35) cells were maintained at 37°C and 5% CO_2_ in Dulbecco’s Modified Eagle Medium High Glucose (DMEM, PAN-Biotech) supplemented with 10% decomplemented Fetal Bovine Serum (FBS, Gibco) and 1% of antibiotics (10 000 units/mL penicillin and 10 000 μg/mL streptomycin, Gibco).

H1N1_WSN_ (strain A/WSN/33 (H1N1)) and H1N1_WSN_ PB2-Nanoluc viruses (36) were provided by the Molecular Genetics of RNA Virus Lab (Pasteur Institute, Paris, France).

### 4.2. GPCA

737 ORFs encoding for 585 UPS factors were assembled in a library by the Molecular Genetics of RNA Virus Lab (Pasteur Institute, Paris, France). This library was cloned through Gateway recombinatorial cloning in the *Gaussia princeps* luciferase protein complementation assay (GPCA) plasmid system, to obtain Glc1-UPS expressing plasmids (33, 37). The ORFs encoding for NS1 from influenza A virus A/WSN/33 (H1N1_WSN_), A/BrevigMission/1/1918 (H1N1_1918_), A/Bretagne/7608/2009 (H1N1_pdm09_), A/Centre/1003/2012 (H3N2), A/Anhui/1/2013 (human H7N9) and A/chicken/Alabama/17– 008901–1/2017 (avian H7N9) strains were amplified by PCR and cloned into Gateway pDONR207. Resulting entry clones were transferred into Gateway pDEST plasmids expressing the Glc2 fragment of *Gaussia princeps* luciferase to obtain NS1-Glc2. Other H1N1_WSN_ proteins (PB1, PB2, PA, NP, M1 and PA-X) were cloned in the same way. The random reference set (RRS) containing human ORFs encoding for random human proteins was cloned into both Glc1 and Glc2 expressing plasmids. Empty Glc1 and empty Glc2 plasmids were used to calculate the normalized luminescence ratio (NLR), taking into account the background luminescence values of each interaction (38).

For the first GPCA screening, the entire UPS library was screened with the 6 NS1-Glc2 coding plasmids and a positive threshold was determined as Q3 + 1.5 IQR for each NS1 screening (Figure S1, Table S1). A GPCA screening validation (all UPS identified in the first screen as interactors of at least 1 NS1) was then performed on positive candidates by using NLR, defined as the fold change normalized over the sum of controls. For a given interacting protein pair A/B, NLR is calculated as (Glc1-A + Glc2-B)/[(Glc1-A + empty Glc2) + (empty Glc1 + Glc2-B)] (37). 20 RRS were also screened against the 6 NS1-Glc2 plasmids and positive threshold was based on the RRS confidence interval of 99.73% (RRS NLR mean + 3 RRS NLR SD) (33) (Table S2). Selected positive interactors were finally screened against H1N1_WSN_ proteins, as well as against 20 RRS during a third GPCA screening, to exclude eventual sticky proteins (Table S3). For each screening, HEK-293T cells were co-transfected with 100 ng of corresponding plasmids using linear PEI (Polyethyleneimine, Polysciences). 24h after transfection, cells were lysed and subjected to luciferase assay, as described in (33).

### 4.3. siRNA assays

Small interfering RNAs were purchased from ThermoFisher Scientific (Validated Silencer Select siRNA, Ambion). For each selected UPS factor, 2 siRNAs were used in each experiment. Three other mRNAs were also targeted by two siRNAs each and served as controls (*ATXN3, RNF40* and *TRAF2*). Finally, one non-targeting siRNA (SCR) was used as a negative depletion control in each experiment (Table S4). For siRNA assays, cells were transfected with 5 nM of siRNA using Lipofectamine RNAiMAX Reagent (ThermoFisher Scientific) for 72 hours.

siRNA efficiency was controlled by RT-qPCR after 72 hours of transfection (Figure S3A). Total RNA extraction was performed on HEK-293T cell lysates using the ReliaPrep RNA Cell Miniprep System (Promega) and a genomic DNA digestion was performed using DNase I (ThermoFisher Scientific) according to the recommendations. From this digestion, RNA was used to perform a reverse-transcription with qScript cDNA Synthesis Kit (QuantaBio). cDNAs were used as templates for qPCR using a LightCycler 480 (Roche) and PerfeCTa SYBR Green SuperMix ROX 2X (QuantaBio), with gene specific forward and reverse primers (Table S5). Cellular GAPDH mRNA served as an internal control.

Cell viability was determined with CellTiter-Glo 2.0 (Promega) in A549 cells, by measuring luminescence after 72 hours of siRNA treatment (Figure S3B).

### 4.4. Viral titration

48 hours after their transfection with siRNA, A549 cells were infected with H1N1_WSN_ at a MOI of 10^−4^ for 24 hours. Plaque assays on MDCK-SIAT cells were performed as described in (39)(Table S6).

For single-cycle infection assays, the H1N1_WSN_ PB2-Nanoluc virus was used to infect siRNA-depleted A549 cells 48 hours post-transfection at a MOI of 10^−4^. After 3 hours, 6 hours, 9 hours, or 12 hours, supernatants containing viral particles were collected and used to infect non-treated A549 cells for 6 hours. The luciferase activity was measured using the Nano-Luciferase assay (Promega) (Table S7).

### 4.5. Minigenome replicon assay

Minigenome replicon assay was performed as in (40) : 48 hours after transfection with siRNA, HEK-293T cells were co-transfected with a combination of plasmids expressing PB1, PB2, PA, NP and NS H1N1_WSN_ viral proteins together with a reporter minigenome pPolI-Firefly (50 ng each plasmid) and 5 ng of pTk-*Renilla* (as transfection efficiency control) plasmids by using JetPEI transfection reagent (Polyplus). 24 hours after plasmid co-transfection, firefly luciferase and *Renilla* luciferase activities were measured using the Dual-Glo luciferase assay system (Promega).

### 4.6. IFN pathway assay

A stable A549 cell line expressing luciferase upon IFN stimulation was previously generated in our lab (34). A549-RING1 cells were transfected with siRNA, and then infected with H1N1_WSN_ at a MOI of 10^−4^ 24 hours post-transfection. Non-infected cells served as controls. 48 hours after infection, firefly luciferase activity was determined by using the One-Glo reagent according to manufacturer’s recommendations (Promega). The results were then normalized to SCR.

### 4.7. Immunofluorescence assay

A549 cells were seeded on coverslips and infected with H1N1_WSN_ virus at a MOI of 3, 24 hours after seeding. After different infection time points (0h, 3h, 6h, 9h and 12h), cells were fixed with PBS-4% paraformaldehyde. Following cell permeabilization with PBS-0.1% Triton X-100 and blocking with PBS-3% BSA, cells were incubated with different mixtures of primary antibodies: FZR1/Cdh1 Rabbit Polyclonal Antibody (16368-1-AP, Proteintech), MKRN3 Rabbit Polyclonal Antibody (PA5-56170, ThermoFisher Scientific), RC3H2 Rabbit Polyclonal Antibody (PA5-60151, ThermoFisher Scientific) or SHKBP1 Rabbit Polyclonal Antibody (PA5-59773, ThermoFisher Scientific), together with Anti-Influenza A Non-Structural Protein 1 (NS1) Mouse Monoclonal Antibody (EMB005, Kerafast). Cells were then washed and incubated with a mixture of secondary antibodies: Fluorescein (FITC)–conjugated Anti-Rabbit IgG (SA00003-2, Proteintech) and Alexa Fluor 568 Goat anti-Mouse IgG (A-11004, ThermoFisher Scientific). Coverslips were mounted in ProLong Gold Antifade Mountant (ThermoFisher Scientific) and analyzed under a confocal microscope (Nikon Eclipse Ti2-E) using x60 oil immersion objective lenses.

### 4.8. Western blot analysis

Cells were lysed in RIPA lysis buffer (50 mM Tris-HCl (pH 7.5), 250 mM NaCl, 2 mM EDTA, 1% Triton-X-100), containing protease inhibitor cocktail (Complete ULTRA Tablets Mini, Roche). After centrifugation of cell lysates, supernatants were collected, and protein extracts were separated by SDS-PAGE and transferred to a PVDF membrane. Protein blots were incubated with primary antibodies overnight at 4°C using the above-mentioned antibodies and the Gaussia Rabbit Polyclonal Antibody (E8023S, New England BioLabs), the Influenza A NS1 Mouse Monoclonal Antibody (sc-130568, Santa Cruz Biotechnology), β-Tubulin Rabbit Polyclonal Antibody (10094-I-AP, Proteintech) and β-Actin Mouse Monoclonal Antibody (sc-69879, Santa Cruz Biotechnology). Blots were incubated with the corresponding secondary antibody for 1 h at room temperature: HRP-conjugated Goat anti-Mouse IgG (G21040, ThermoFischer Scientific), HRP-conjugated Goat anti-Rabbit IgG (SA00001-2, Proteintech). ECL Western Blotting Substrate (ThermoFischer Scientific) was used as enhanced chemiluminescence substrate. ImageJ was used to quantify protein abundance by densitometry (41).

## Acknowledgements

We thank the UMONS Cell Biology lab members for their helpful discussions, and particularly E. Hennebert for the confocal microscopy training. We are grateful to the Molecular Genetics of RNA Virus Lab (Pasteur Institute, Paris, France) for providing both H1N1_WSN_ and H1N1_WSN_ PB2-Nanoluc viruses, and to Y. Jacob for Nano-luciferase luminescence protocol. We also thank M. Matrosovich for providing MDCK-SIAT cells, Metabolic and Molecular Biochemistry lab for LightCycler 480 use, and C. Barbezange (Sciensano, Brussels, Belgium) for technical help in IAV amplification and titration assay. L.-A.L. is supported by a FRIA PhD fellowship grant 5117919F. This work was supported by the Fonds de la Recherche Scientifique-FNRS under CDR grants J.0198.20 and J.0172.23, and by the Fonds pour la Recherche Médicale dans le Hainaut-FRMH. The funders had no role in study design, data collection and interpretation, or the decision to submit the work for publication.

## Supplementary figure legends

**Figure S1: GPCA screening with 6 NS1 against the human UPS library.** Whisker-plots were generated from luminescence values (RLU) of the 737 candidates and the positive threshold was determined as Q3 + 1.5 IQR for each NS1. Outlier luminescence values of tested factors are considered as positive candidates (circles for NS1 of H1N1_WSN_, squares for H1N1_1918_, up triangles for H1N1_pdm09_, down triangles for H3N2, diamonds for H7N9 and crosses for avian H7N9) and few examples are identified.

**Figure S2: Assessment of the different NS1-Gluc2 expression level for GPCA.** After GPCA luminescence measurements, protein extracts of cells transfected by pGluc1 and pGluc2-NS1 (from the 6 different strains) were subjected to western-blot. Anti-Gaussia antibody revealed the NS1-Gluc2 fusion proteins, anti-tubulin was used as loading control.

**Figure S3: Silencing efficiency and toxicity of siRNAs.** (A) HEK-293T cells were transfected with 2 siRNAs/factor for 72h. Total RNA was extracted, followed by a genomic DNA digestion and RT-PCR were performed on each RNA extract. cDNAs were used as templates for qPCR using gene specific forward and reverse primers. Cellular GAPDH mRNA served as an internal control. Results represent the percentage of remaining mRNA in each condition, as compared to SCR (calculated using the ΔΔCt ratio). No amplification was possible for ZBTB32 gene (n.a.). ATXN3, RNF40 and TRFA2, in bold, are literature controls. qPCR were performed in triplicates and are represented as the mean of triplicate. (B) A549 cell viability was assessed upon siRNA-mediated depletion of UPS factors (2 siRNAs/factor) for 72h by using CellTiter-Glo 2.0 (Promega). Luminescence values for each condition were normalized to SCR-transfected cells. Data represent means ± SD of three independent experiments, and p values were calculated with unpaired t-tests, *<0.05, **<0.01, ***<0.001.

**Figure S4: Standardization of plaque and Nano-luciferase assays.** A549 cells were infected with the H1N1_WSN_ PB2-Nanoluc virus at a MOI of 10^−4^. At 24 hours post-infection, the supernatant was recovered and diluted (dilutions 5×10^4^, 1×10^5^, 5×10^5^ and 1×10^6^). Each dilution was used to either perform a plaque titration assay or a Nano-luciferase titration assay. Results are expressed as number of lysis plaques for plaque assay (purple) and relative luminescence units (RLU) for Nano-luciferase assay (yellow). Data represent the mean ± SD of ten independent experiments.

**Figure S5: Effects of H1N1_WSN_ infection on UPS factor transcription.** A549 cells were infected with H1N1_WSN_ at a MOI of 10^−4^. At 48 hours post-infection, cells were lysed and a total RNA extraction, followed by genomic DNA digestion and RT-PCR, was performed. cDNAs from infected and from mock-infected cells were used as templates for qPCR, using factors specific forward and reverse primers. Cellular GAPDH served as an internal control. Results are expressed as the percentage of mRNA of each factor in infected cells, as compared to mock-infected cells (100%). No cDNA amplification was possible for BCL6B, DZIP3 and TRIM4 conditions (n.a.). ATXN3, RNF40 and TRFA2, in bold, are literature controls. RIG-I and MDA5 (orange) are interferon-stimulated factors. qPCR were performed in triplicates and are represented as the mean of triplicate.

## References

1. Dikic I, Dötsch V. 2009. Ubiquitin linkages make a difference. Nat Struct Mol Biology 16:1209–1210.

2. Chen ZJ, Sun LJ. 2009. Nonproteolytic Functions of Ubiquitin in Cell Signaling. Mol Cell 33:275–286.

3. Chau V, Tobias JW, Bachmair A, Marriott D, Ecker DJ, Gonda DK, Varshavsky A. 1989. A Multiubiquitin Chain Is Confined to Specific Lysine in a Targeted Short-Lived Protein. Science 243:1576–1583.

4. Ciechanover A. 2015. The unravelling of the ubiquitin system. Nat Rev Mol Cell Bio 16:322–324.

5. Hershko A, Heller H, Elias S, Ciechanover A. 1983. Components of ubiquitin-protein ligase system. Resolution, affinity purification, and role in protein breakdown. J Biological Chem 258:8206–8214.

6. Veen AG van der, Ploegh HL. 2012. Ubiquitin-Like Proteins. Annu Rev Biochem 81:323– 357.

7. Komander D, Rape M. 2012. The Ubiquitin Code. Annu Rev Biochem 81:203–229.

8. Kulathu Y, Komander D. 2012. Atypical ubiquitylation — the unexplored world of polyubiquitin beyond Lys48 and Lys63 linkages. Nat Rev Mol Cell Biology 13:508–523.

9. Tanno H, Komada M. 2013. The ubiquitin code and its decoding machinery in the endocytic pathway. The J Biochem 153:497–504.

10. Komander D, Clague MJ, Urbé S. 2009. Breaking the chains: structure and function of the deubiquitinases. Nat Rev Mol Cell Biology 10:550–563.

11. Clague MJ, Barsukov I, Coulson JM, Liu H, Rigden DJ, Urbé S. 2013. Deubiquitylases From Genes to Organism. Physiol Rev 93:1289–1315.

12. Gent M van, Sparrer KMJ, Gack MU. 2018. TRIM Proteins and Their Roles in Antiviral Host Defenses. Annu Rev Virol 5:385–405.

13. Lee H-R, Lee MK, Kim CW, Kim M. 2020. TRIM Proteins and Their Roles in the Influenza Virus Life Cycle. Microorganisms 8:1424.

14. Gack MU, Albrecht RA, Urano T, Inn K-S, Huang I-C, Carnero E, Farzan M, Inoue S, Jung JU, García-Sastre A. 2009. Influenza A virus NS1 targets the ubiquitin ligase TRIM25 to evade recognition by the host viral RNA sensor RIG-I. Cell host & microbe 5:439–449.

15. Gack MU, Shin YC, Joo C-H, Urano T, Liang C, Sun L, Takeuchi O, Akira S, Chen Z, Inoue S, Jung JU. 2007. TRIM25 RING-finger E3 ubiquitin ligase is essential for RIG-I-mediated antiviral activity. Nature 446:916–920.

16. Krammer F, Smith GJD, Fouchier RAM, Peiris M, Kedzierska K, Doherty PC, Palese P, Shaw ML, Treanor J, Webster RG, García-Sastre A. 2018. Influenza. Nature reviews Disease primers 4:3–21.

17. Talon J, Horvath CM, Polley R, Basler CF, Muster T, Palese P, García-Sastre A. 2000. Activation of Interferon Regulatory Factor 3 Is Inhibited by the Influenza A Virus NS1 Protein. J Virol 74:7989–7996.

18. Rückle A, Haasbach E, Julkunen I, Planz O, Ehrhardt C, Ludwig S. 2012. The NS1 Protein of Influenza A Virus Blocks RIG-I-Mediated Activation of the Noncanonical NF-κB Pathway and p52/RelB-Dependent Gene Expression in Lung Epithelial Cells. J Virol 86:10211–10217.

19. Wang X, Li M, Zheng H, Muster T, Palese P, Beg AA, Garcıa-Sastre A. 2000. Influenza A Virus NS1 Protein Prevents Activation of NF-κB and Induction of Alpha/Beta Interferon. J Virol 74:11566–11573.

20. Min J-Y, Li S, Sen GC, Krug RM. 2007. A site on the influenza A virus NS1 protein mediates both inhibition of PKR activation and temporal regulation of viral RNA synthesis. Virology 363:236–243.

21. Min J-Y, Krug RM. 2006. The primary function of RNA binding by the influenza A virus NS1 protein in infected cells: Inhibiting the 2′-5′ oligo (A) synthetase/RNase L pathway. Proc National Acad Sci 103:7100–7105.

22. Marc D. 2014. Influenza virus non-structural protein NS1: interferon antagonism and beyond. The Journal of general virology 95:2594–2611.

23. Carrillo B, Choi J-M, Bornholdt ZA, Sankaran B, Rice AP, Prasad BVV. 2014. The Influenza A Virus Protein NS1 Displays Structural Polymorphism. J Virol 88:4113–4122.

24. Ludwig S, Wang X, Ehrhardt C, Zheng H, Donelan N, Planz O, Pleschka S, García-Sastre A, Heins G, Wolff T. 2002. The Influenza A Virus NS1 Protein Inhibits Activation of Jun N-Terminal Kinase and AP-1 Transcription Factors. J Virol 76:11166–11171.

25. Mibayashi M, Martínez-Sobrido L, Loo Y-M, Cárdenas WB, Gale M, García-Sastre A. 2007. Inhibition of retinoic acid-inducible gene I-mediated induction of beta interferon by the NS1 protein of influenza A virus. Journal of Virology 81:514–524.

26. Pizzorno A, Dubois J, Machado D, Cartet G, Traversier A, Julien T, Lina B, Bourdon J-C, Rosa-Calatrava M, Terrier O. 2018. Influenza A viruses alter the stability and antiviral contribution of host E3-ubiquitin ligase Mdm2 during the time-course of infection. Scientific reports 8:3746.

27. Jahan AS, Biquand E, Muñoz-Moreno R, Quang AL, Mok CK-P, Wong HH, Teo QW, Valkenburg SA, Chin AWH, Poon LLM, Velthuis AT, García-Sastre A, Demeret C, Sanyal S. 2020. OTUB1 Is a Key Regulator of RIG-I-Dependent Immune Signaling and Is Targeted for Proteasomal Degradation by Influenza A NS1. Cell reports 30:1570–1584.e6.

28. Maelfait J, Roose K, Vereecke L, Guire CM, Sze M, Schuijs MJ, Willart M, Ibañez LI, Hammad H, Lambrecht BN, Beyaert R, Saelens X, Loo G van. 2016. A20 Deficiency in Lung Epithelial Cells Protects against Influenza A Virus Infection. PLoS Pathog 12:e1005410.

29. Feng W, Sun X, Shi N, Zhang M, Guan Z, Duan M. 2017. Influenza a virus NS1 protein induced A20 contributes to viral replication by suppressing interferon-induced antiviral response. Biochemical and biophysical research communications 482:1107–1113.

30. Lamotte L-A, Tafforeau L. 2021. How Influenza A Virus NS1 Deals with the Ubiquitin System to Evade Innate Immunity. Viruses 13:2309.

31. Muller M, Jacob Y, Jones L, Weiss A, Brino L, Chantier T, Lotteau V, Favre M, Demeret C. 2012. Large scale genotype comparison of human papillomavirus E2-host interaction networks provides new insights for e2 molecular functions. PLoS pathogens 8:e1002761.

32. Poirson J, Biquand E, Straub M-L, Cassonnet P, Nominé Y, Jones L, Werf S van der, Travé G, Zanier K, Jacob Y, Demeret C, Masson M. 2017. Mapping the interactome of HPV E6 and E7 oncoproteins with the ubiquitin-proteasome system. The FEBS journal 284:3171–3201.

33. Biquand E, Poirson J, Karim M, Declercq M, Malausse N, Cassonnet P, Barbezange C, Straub M-L, Jones L, Munier S, Naffakh N, Werf S van der, Jacob Y, Masson M, Demeret C. 2017. Comparative Profiling of Ubiquitin Proteasome System Interplay with Influenza A Virus PB2 Polymerase Protein Recapitulating Virus Evolution in Humans. Msphere 2:e00330–17.

34. Lamotte L-A, Tafforeau L. 2023. Generation of an A549 ISRE-luciferase stable cell line. J Virol Methods 316:114731.

35. Matrosovich M, Matrosovich T, Carr J, Roberts NA, Klenk H-D. 2003. Overexpression of the alpha-2,6-sialyltransferase in MDCK cells increases influenza virus sensitivity to neuraminidase inhibitors. Journal of Virology 77:8418–8425.

36. Diot C, Fournier G, Santos MD, Magnus J, Komarova A, Werf S van der, Munier S, Naffakh N. 2016. Influenza A Virus Polymerase Recruits the RNA Helicase DDX19 to Promote the Nuclear Export of Viral mRNAs. Sci Rep-uk 6:33763.

37. Cassonnet P, Rolloy C, Neveu G, Vidalain P-O, Chantier T, Pellet J, Jones L, Muller M, Demeret C, Gaud G, Vuillier F, Lotteau V, Tangy F, Favre M, Jacob Y. 2011. Benchmarking a luciferase complementation assay for detecting protein complexes. Nature Methods 8:990– 992.

38. Neveu G, Cassonnet P, Vidalain P-O, Rolloy C, Mendoza J, Jones L, Tangy F, Muller M, Demeret C, Tafforeau L, Lotteau V, Rabourdin-Combe C, Travé G, Dricot A, Hill DE, Vidal M, Favre M, Jacob Y. 2012. Comparative analysis of virus-host interactomes with a mammalian high-throughput protein complementation assay based on Gaussia princeps luciferase. Methods (San Diego, Calif) 58:349–359.

39. Matrosovich M, Matrosovich T, Garten W, Klenk H-D. 2006. New low-viscosity overlay medium for viral plaque assays. Virol J 3:63.

40. Tafforeau L, Chantier T, Pradezynski F, Pellet J, Mangeot PE, Vidalain P-O, André P, Rabourdin-Combe C, Lotteau V. 2011. Generation and comprehensive analysis of an influenza virus polymerase cellular interaction network. Journal of Virology 85:13010–13018.

41. Schneider CA, Rasband WS, Eliceiri KW. 2012. NIH Image to ImageJ: 25 years of image analysis. Nat Methods 9:671–675.

42. Li Y, Yamakita Y, Krug RM. 1998. Regulation of a nuclear export signal by an adjacent inhibitory sequence: The effector domain of the influenza virus NS1 protein. Proc National Acad Sci 95:4864–4869.

43. Melén K, Kinnunen L, Fagerlund R, Ikonen N, Twu KY, Krug RM, Julkunen I. 2007. Nuclear and nucleolar targeting of influenza A virus NS1 protein: striking differences between different virus subtypes. Journal of Virology 81:5995–6006.

44. Volmer R, Mazel-Sanchez B, Volmer C, Soubies SM, Guérin J-L. 2010. Nucleolar localization of influenza A NS1: striking differences between mammalian and avian cells. Virol J 7:63.

45. Luo H. 2016. Interplay between the virus and the ubiquitin-proteasome system: molecular mechanism of viral pathogenesis. Current opinion in virology 17:1–10.

46. Li L, Gong Y, Tang J, Yan C, Li L, Peng W, Cheng Z, Yu R, Xiang Q, Deng C, Mu J, Xia J, Luo X, Wu Y, Xiang T. 2022. ZBTB28 inhibits breast cancer by activating IFNAR and dual blocking CD24 and CD47 to enhance macrophages phagocytosis. Cell Mol Life Sci 79:83.

47. Xu D, Holko M, Sadler AJ, Scott B, Higashiyama S, Berkofsky-Fessler W, McConnell MJ, Pandolfi PP, Licht JD, Williams BRG. 2009. Promyelocytic Leukemia Zinc Finger Protein Regulates Interferon-Mediated Innate Immunity. Immunity 30:802–816.

48. Maruyama T, Araki T, Kawarazaki Y, Naguro I, Heynen S, Aza-Blanc P, Ronai Z, Matsuzawa A, Ichijo H. 2014. Roquin-2 Promotes Ubiquitin-Mediated Degradation of ASK1 to Regulate Stress Responses. Sci Signal 7:ra8.

49. Yang Y, Bazhin AV, Werner J, Karakhanova S. 2013. Reactive Oxygen Species in the Immune System. Int Rev Immunol 32:249–270.

50. Chen I-Y, Ichinohe T. 2015. Response of host inflammasomes to viral infection. Trends Microbiol 23:55–63.

51. Garzia A, Jafarnejad SM, Meyer C, Chapat C, Gogakos T, Morozov P, Amiri M, Shapiro M, Molina H, Tuschl T, Sonenberg N. 2017. The E3 ubiquitin ligase and RNA-binding protein ZNF598 orchestrates ribosome quality control of premature polyadenylated mRNAs. Nat Commun 8:16056.

52. DiGiuseppe S, Rollins MG, Bartom ET, Walsh D. 2018. ZNF598 Plays Distinct Roles in Interferon-Stimulated Gene Expression and Poxvirus Protein Synthesis. Cell Rep 23:1249– 1258.

53. Wang G, Kouwaki T, Okamoto M, Oshiumi H. 2019. Attenuation of the Innate Immune Response against Viral Infection Due to ZNF598-Promoted Binding of FAT10 to RIG-I. Cell reports 28:1961–1970.e4.

54. Feng L, Wang J-T, Jin H, Qian K, Geng J-G. 2011. SH3KBP1-binding protein 1 prevents epidermal growth factor receptor degradation by the interruption of c-Cbl-CIN85 complex. Cell Biochem Funct 29:589–96.

55. Eierhoff T, Hrincius ER, Rescher U, Ludwig S, Ehrhardt C. 2010. The Epidermal Growth Factor Receptor (EGFR) Promotes Uptake of Influenza A Viruses (IAV) into Host Cells. PLoS Pathog 6:e1001099.

56. Wee P, Wang Z. 2017. Epidermal Growth Factor Receptor Cell Proliferation Signaling Pathways. Cancers 9:52.

57. Zhou Y, Ching Y-P, Ng RWM, Jin D-Y. 2003. Differential expression, localization and activity of two alternatively spliced isoforms of human APC regulator CDH1. Biochem J 374:349–358.

58. Zhou Y, Ching Y-P, Chun ACS, Jin D-Y. 2003. Nuclear Localization of the Cell Cycle Regulator CDH1 and Its Regulation by Phosphorylation*. J Biol Chem 278:12530–12536.

59. He Y, Xu K, Keiner B, Zhou J, Czudai V, Li T, Chen Z, Liu J, Klenk H-D, Shu YL, Sun B. 2010. Influenza A Virus Replication Induces Cell Cycle Arrest in G 0 /G 1 Phase. J Virol 84:12832– 12840.

60. Jiang W, Wang Q, Chen S, Gao S, Song L, Liu P, Huang W. 2013. Influenza A Virus NS1 Induces G 0 /G 1 Cell Cycle Arrest by Inhibiting the Expression and Activity of RhoA Protein. J Virol 87:3039–3052.

61. Nishimura K, Oki T, Kitaura J, Kuninaka S, Saya H, Sakaue-Sawano A, Miyawaki A, Kitamura T. 2013. APC(CDH1) targets MgcRacGAP for destruction in the late M phase. Plos One 8:e63001.

62. Kolapalli SP, Sahu R, Chauhan NR, Jena KK, Mehto S, Das SK, Jain A, Rout M, Dash R, Swain RK, Lee DY, Rusten TE, Chauhan S, Chauhan S. 2021. RNA-Binding RING E3-Ligase DZIP3/hRUL138 Stabilizes Cyclin D1 to Drive Cell-Cycle and Cancer Progression. Cancer Res 81:315–331.

63. Zhou J, Xu Y, Lin S, Guo Y, Deng W, Zhang Y, Guo A, Xue Y. 2018. iUUCD 2.0: an update with rich annotations for ubiquitin and ubiquitin-like conjugations. Nucleic Acids Res 46:D447–D453.

